# Cryo-EM Structures of Infectious Low-Twist α-Synuclein Filaments in Lewy Bodies

**DOI:** 10.64898/2026.09.16.752119

**Authors:** Nicholas L. Yan, Mylan Mayer, Arthur A. Melo, Jacob I. Ayers, Daniel A. Mordes, William W. Seeley, Stanley B. Prusiner, Eric Tse, Vinita G. Chittoor-Vinod, Gregory E. Merz

## Abstract

α-Synuclein (aSyn) aggregation within Lewy bodies underlies Parkinson’s disease and associated dementias. These aggregates are often comprised of amyloid filaments, most of which appear untwisted or have extremely low twist, complicating high-resolution structural determination by cryogenic electron microscopy (cryo-EM). Here, we isolated filaments from the brain tissue of a patient with dementia with Lewy bodies and show infectivity in a cell line system that propagates Lewy aSyn. Then, we determined 2.3–2.9 Å cryo-EM structures of low-twist aSyn filaments from this tissue and from a publicly available Parkinson’s disease dementia dataset. In both structures, the protofilament adopts a fold highly similar to the canonical Lewy fold with conserved cofactor and peptidic densities. Compared to previously resolved high-twist filaments, the low-twist filament is left-handed and flattened along the filament axis, resulting in register shifts of interlayer contacts, while preserving the overall architecture. Our results establish that low-twist filaments, which include over 75% of disease-associated Lewy aSyn amyloids, share a common structural core with high-twist forms and are the predominant aSyn species in samples that engage in templated infection.

## INTRODUCTION

Parkinson’s disease (PD), Parkinson’s disease dementia (PDD), and dementia with Lewy bodies (DLB) are clinical syndromes that result from underlying Lewy body disease (LBD), the neuropathological entity defined by accumulation of intracellular inclusions known as Lewy bodies and Lewy neurites[29]. PDD and DLB are collectively referred to as Lewy body dementias (LBD), a group of clinically overlapping disorders distinguished from other dementias by the presence of Lewy body (LB) pathology.

These pathological aggregates are composed primarily of misfolded α-synuclein (aSyn), a 140-residue, intrinsically disordered presynaptic protein that is abundant in neurons[7, 8]. The progressive conversion of soluble aSyn into β-sheet-rich amyloid filaments via templating and propagation is a hallmark of diseases known as synucleinopathies and is thought to play a central role in disease pathogenesis, contributing to synaptic dysfunction, neuronal loss, and pathological spread through connected brain regions[3, 22]. Elucidating the molecular architecture of aSyn filaments is therefore critical for understanding disease etiology, as filament structure is expected to encode key pathogenic properties, including aggregation kinetics, seeding capacity, cellular interactions, and strain-specific differences that may underlie clinical heterogeneity across synucleinopathies[12]. Despite extensive study, the precise structures of aSyn filaments within Lewy bodies remain incompletely understood.

Advances in cryogenic electron microscopy (cryo-EM) have enabled the elucidation of near-atomic resolution structures of amyloid filaments extracted directly from the brain tissue of patients with synucleinopathies. These structures reveal that disease-associated aggregates can adopt distinct, disease-specific folds, correlating with differences in clinical presentation, progression, and cellular vulnerability[11, 27, 28]. For aSyn, several filament structures have been reported from *in vitro* assemblies and *ex vivo* from synucleinopathies such as multiple system atrophy (MSA) and juvenile-onset synucleinopathy[10, 26, 31, 32]. In 2022, the first high-resolution structures derived from aSyn filaments isolated from the brain tissue of patients with PD, PDD, and DLB were reported, revealing right-handed helical filaments with a shared fold that is largely distinct from previously described structures[33]. The ordered core of this “Lewy fold” consists of a single protofilament (PF) spanning residues 31–100. The Lewy fold adopts a 3-layered fold comprising a Lys-rich N-terminal cavity (residues 31–45), a central hinge region (46–68), and a C-terminal hairpin (72–100). Additional unidentified densities are also present, including one within the N-terminal cavity and two peptidic “islands”: island A, which associates with the hinge region, and island B, which binds to the C-terminal hairpin.

Although these structures represent a major advance, structural characterization is limited by the fact that only a minority of filament segments (up to ∼25%) are amenable to high-resolution structural determination due to their well-defined twist. Most segments appear untwisted in 2D classification. This hinders helical reconstruction attempts due to the potential for preferred orientation and lower signal-to-noise of these particles, compared to twisted filaments that adopt a full range of views around the filament axis[19]. To address this limitation, a 2024 study reported solid-state nuclear magnetic resonance structures of serially amplified low-twist aSyn filaments seeded from LBD tissue, revealing shared motifs with the Lewy fold (although not full recapitulation)[9]. However, this approach requires milligram quantities of labeled material, limiting its applicability in primary tissue samples. With the fold of the untwisted LB filament segments yet unknown, rational design of diagnostics and therapeutics for these diseases is done at risk. Differential twisted and untwisted folds may cause small molecules to bind only to a subset of filament types, limiting their clinical application. Therefore, understanding the structure of the untwisted and low-twist filaments is crucial in both disease biology and medicinal chemistry contexts.

To more fully characterize the low-twist aSyn filaments that comprise the majority of those found in the LBs, we isolated filaments from brain tissue of a patient with DLB and validated their infectivity in a stable cell line expressing a fluorescently tagged aSyn construct that propagates prions with the Lewy fold[4]. Then, using recent advances in cryo-EM image processing, notably Blush regularization[17, 20], we determined 2.3–2.9 Å cryo-EM structures of low-twist aSyn filaments derived from the DLB tissue and from a publicly available PDD dataset. In both cases, aSyn adopted the Lewy fold with a conserved pattern of flanking peptidic and non-peptidic densities. However, relative to high-twist filaments, the low-twist PF was “flatter” along the filament axis, resulting in shifts in otherwise shared intrafilament contacts.

## METHODS

### aSyn filament extraction

Sarkosyl-insoluble material was extracted from freshly frozen inferior temporal gyrus of a 74-year-old female patient with DLB using a protocol similar to previous aSyn filament purifications[26] (Supplementary Fig. S1). The same extraction protocol was applied to tissue from the anterior internal capsule of a 78-year-old female patient with MSA.

Tissue was homogenized in 20 volumes (v/w) of extraction buffer composed of 10 mM Tris-HCl, pH 7.5, 0.8 M NaCl, 10% sucrose, and 1 mM ethyleneglycol- bis(β-aminoethyl)-N,N,N ,N -tetraacetic acid (EGTA). Fraction 1 was collected from this homogenate; the remaining homogenate was brought to 2% sarkosyl, incubated at 37 °C for 30 min, and then centrifuged at 10,000 × g for 10 min. The supernatants were then collected and pooled, and the pellets were resuspended in 20 volumes (v/w) of extraction buffer with 2% sarkosyl. The resuspended pellets were centrifuged at 10,000 × g for another 10 min, and the supernatants were again collected and added to the pooled supernatants from the previous step. Fraction 2 was collected from the supernatant pool, and the remaining solution was divided and spun at 100,000 × g for 30 min at 22 °C. The pellets were then isolated, resuspended in 500 μL/g extraction buffer, and combined. Fraction 3 was collected from the resuspension solution. The remaining solution was then centrifuged at 3,000 × g for 5 min. The new supernatants were combined and diluted 3-fold in 50 mM Tris-HCl, pH 7.5, 0.15 M NaCl, 10% sucrose, and 0.2% sarkosyl, and fraction 4 was collected from the diluted solution. The remaining solution was centrifuged at 166,000 × g for 40 min at 22 °C. The final sarkosyl-insoluble pellet was resuspended in 100 μL/g of 50 mM Tris-HCl, pH 7.5. The resuspended pellet was centrifuged at 3,000 × g for 5 min to separate any insoluble debris, and the supernatant was used for negative stain and cryo-EM experiments. The final supernatant was designated as fraction 5.

### Negative-stain electron microscopy

The sarkosyl-insoluble preparation from DLB tissue was diluted 1:5 and 1:10, and 5 μL of sample was added to a glow-discharged formvar-coated 300 mesh copper grid covered with a layer of amorphous carbon. After approximately 30 s, the grid was blotted with filter paper, then 5 μL of water was added and blotted with filter paper to wash. The wash step was repeated once more for a total of 2 washes. Then, 5 μL of 0.75% uranyl formate was added and blotted with filter paper after approximately 40 s. Another 5 μL of uranyl formate was added and then removed by gradual vacuum aspiration; this was repeated once more for a total of 3 aliquots of uranyl formate. Images were collected on a Talos L120C transmission electron microscope (Thermo Fisher Scientific) operating at 120 kV and equipped with a Ceta-D camera (Thermo Fisher Scientific).

### Cell lines and infectivity assays

HEK293T cell lines stably expressing aSyn(35–110)*E46K-YFP or aSyn(35–110)*A53T-YFP were generated in house using previously described methods[30]. All cells were cultured in 1X Dulbecco’s Modified Eagle’s Medium (Corning), supplemented with 10% (v/v) fetal bovine serum (Avantor, Inc.) and 50 U/mL penicillin/streptomycin (Gibco).

These cells were maintained in a humidified environment at 37 °C and 5% CO_2_. For infectivity assays, cells were plated at 2,800 or 4,000 cells/well in a 384-well plate with black polystyrene walls (Greiner Bio-One). Hoechst 33342 (Invitrogen) was added to the cells at 0.1 μg/mL to stain the nuclei. Cells were allowed to rest in the incubator for 5 h before transfection. Total protein concentrations of brain fractions were determined by bicinchoninic acid (BCA) assay using the manufacturer’s protocol (Pierce). Fractions were diluted in Dulbecco’s phosphate-buffered saline (Gibco) to achieve the final protein concentrations. Transfection complexes were prepared by mixing the diluted fractions with Lipofectamine-2000 (Thermo Fisher Scientific) in Opti-MEM reduced serum medium (Gibco) and incubating for 2.5 or 1.5 h at room temperature. After incubation time, transfection complexes were added to plated cells in 4 replicate wells and returned to the incubator for 4 d. Individual plates were then imaged on the Opera Phenix Plus High-Content Imaging System (Revvity, Inc.). Images in the 4′,6-diamidino-2-phenylindole (DAPI) and fluorescein isothiocyanate (FITC) channels were collected at 5 predetermined regions within each well. Raw images were analyzed using Harmony High-Content Imaging and Analysis software, using an analysis protocol designed to detect and quantify intracellular aggregates and nuclei numbers (Revvity, Inc.). To evaluate infectivity, CWA% (percent cells with aggregates) was calculated and plotted as mean ± standard deviation (SD) using GraphPad Prism statistical software v11.0.2. Technical replicates were averaged across 4 wells. Infectivity between fraction 5 and other fractions of the same brain sample were compared using unpaired Student’s *t*-tests. The same methods were used to assess infectivity between fractions 5 of DLB and MSA samples.

### Cryo-EM sample preparation and data collection

For the DLB sample, purified filaments (3 μL) (Supplementary Fig. S2) were added to a 200 mesh 1.2/1.3R Au Quantifoil grid coated with a 2-nm thick carbon layer, which was not glow discharged. After 30 s, grids were blotted for 7.5 s at room temperature and 100% humidity using a FEI Vitrobot Mark IV (Thermo Fisher Scientific), followed by plunge freezing in liquid ethane. A total of 88,487 movies over 5 sessions were collected at a nominal magnification of 165,000× (physical pixel size: 0.728 Å/pixel) on a Titan Krios G4 cryogenic transmission electron microscope (Thermo Fisher Scientific) operated at 300 kV and equipped with a Falcon 4i direct electron detector and Selectris X energy filter (Thermo Fisher Scientific) set to a slit width of 20 eV. A defocus range of −0.8 to −1 μm was used with a total exposure time of 4.5–5 s fractionated into 90 TIFF frames. The total dose for each movie was 45 electrons/Å^2^. An additional dataset of 28,158 movies was collected at a 20° tilt, for a total of 116,645 movies. The movies were motion corrected using MotionCor2[35] in Scipion[6]. Motion-corrected and dose-weighted micrographs were manually curated in Scipion to remove micrographs that lacked filaments, were at low resolution, or had significant ice contamination, resulting in 6,623 remaining micrographs (4,826 non-tilted and 1,797 tilted data).

### Cryo-EM image processing

#### DLB sample

A graphical overview of the data processing workflow is provided in Supplementary Figure S3. All image processing was done in RELION v4, v5, or v5.1[13, 16, 20]. Dose-weighted summed micrographs were imported into RELION 4. The contrast transfer function was estimated using CTFFIND-4.1[24]. Filaments were manually picked, and segments were extracted with a box size of 900 pixels downscaled to 300 pixels, resulting in 459,285 segments (337,269 non-tilted and 122,016 tilted data). Reference-free 2D classification was used to remove contaminants, non-aSyn filaments, and low-resolution classes, resulting in 331,207 remaining segments (232,804 non-tilted and 98,403 tilted data) (Supplementary Figs. S4–S5). These were re-extracted with a box size of 288 pixels without downscaling for subsequent 3D classification.

In the non-tilted datasets, 2D class averages that displayed a twist (47,500 particles) were used to generate an *ab initio* 3D map using the relion_helix_inimodel2d feature. Multiple crossovers from 800–2,000 Å were tested, and a crossover of 800 Å gave the best alignment from examination of the projections.spi file. An expanded subset of 73,549 segments displaying any twist was used for an initial round of 3D classification with a 15-Å low-pass filtered *ab initio* map as the reference, with k = 4. A regularization parameter (T) of 10 was used for 25 iterations, which was increased to 20 for an additional 7 iterations. The angular sampling interval was set at 1.8°. The tube diameter was set at −1 inner and 175 Å outer. The number of unique asymmetrical units was set to 3. The initial twist and rise were set to −1.07° and 4.76 Å, respectively. Local symmetry searches were enabled as followed: min twist = −1.1°, max twist = −0.2°, step = 0.01°; min rise = 4.7 Å, max rise = 4.8 Å, step = 0.005 Å. All other 3D classification parameters were RELION defaults. After 32 iterations, one class (47,402 segments) displayed connected density that resembled the C-terminal hairpin and island B, although the remainder of the filament was not immediately discernable. This class had a helical twist of −0.43° and a helical rise of 4.78 Å. Due to the relatively small number of segments in this class, we did not pursue additional 3D classification on this particle stack, but instead used a 15-Å low-pass filtered version of this map for 3D classification involving all 232,804 segments in the non-tilted datasets, including those in non-twisting 2D averages. The parameters were as follows: k = 8, T = 10, 35 iterations, initial twist = −1.0°, initial rise = 4.76 Å, min twist = −1.1°, max twist = 0°, twist step = 0.01°; min rise = 4.7 Å, max rise = 4.8 Å, rise step = 0.005 Å, with other parameters being identical to the first round of 3D classification. Afterwards, one class (145,070 segments) displayed backbone density for the C-terminal hairpin and island B. In addition, the class also had features of the central hinge, N-terminal pocket, and island A, although these were less well resolved.

This class had a helical twist of −0.13° and a helical rise of 4.80 Å. This particle stack was subjected to another round of 3D classification (52 iterations) with the following parameters: k = 5, T = 10, initial twist = −0.1°, initial rise = 4.76 Å, min twist = −1°, max twist = 0°, twist step = 0.01°; min rise = 4.7 Å, max rise = 4.9 Å, rise step = 0.005 Å, with other parameters unchanged. The resulting class (72,791 segments) had improved resolution and backbone connectivity for the entire protofilament but weak sidechain density. This refined particle stack was combined with the tilted dataset (98,403 segments; total 171,374 segments) and subjected to 2 more rounds of 3D classification. The parameters for the first round were k = 6, T = 10, 45 iterations, initial twist = −0.2°, with other parameters unchanged, giving a class with 107,312 segments. This class was used for the final round of 3D classification with the parameters k = 4, T = 10, 55 iterations, initial twist = −0.1°, initial rise = 4.8 Å, and all other parameters unchanged. The best class, consisting of 60,550 segments, underwent 3D auto-refinement in RELION v5.1 using Blush regularization with the AmyBlush (amy-v1.0) network[20] and the following parameters: initial twist = −0.1°, initial rise = 4.8 Å, central Z length = 10%, min twist = −0.2°, max twist = 0°, twist step = 0.001°; min rise = 4.7 Å, max rise = 4.9 Å, rise step = 0.001 Å. The refined map then underwent mask creation (initial binarization threshold = 0.001, binary map extension and soft edge = 10 pixels, central Z length = 20%) and standard post-processing to give the final DLB Lewy aSyn map at an estimated global resolution of 2.9 Å (Supplementary Fig. S6). We attempted further processing with contrast transfer function (CTF) refinement, but this did not improve the map quality. We also attempted to resolve short-crossover filaments from this dataset but were unsuccessful. Finally, only twist parameters with a narrow negative search range gave meaningful high-resolution maps.

#### PDD sample

A graphical overview of the data processing workflow is provided in Supplementary Figure S7. All image processing was done in RELION v4, v5, or v5.1. Movies (10,872) downloaded from EMPIAR-11140 were imported into RELION 4. Motion correction was done using RELION’s implementation of MotionCor2. The contrast transfer function was estimated using CTFFIND-4. From the provided start-end coordinates in the data, segments were extracted with a box size of 900 pixels downscaled to 300 pixels, resulting in 1,339,205 segments. Reference-free 2D classification was used to remove contaminants, non-aSyn filaments, and low-resolution classes, resulting in 725,340 remaining segments that had both non-twisted and slightly twisted aSyn (Supplementary Fig. S8). These were re-extracted with a box size of 288 pixels without downscaling for subsequent 3D classification. The unbinned segments underwent one round of 3D classification using a 15-Å low-pass filtered DLB map as a reference and Blush regularization using the default v1.0 network in RELION v5. The other parameters were as follows: 16 iterations, k = 8, T = 10, angular sampling interval = 1.8°, outer tube diameter = 175 Å, number of unique asymmetrical units = 3, initial twist = −0.11°, initial rise = 4.8 Å, central Z length = 10%, no local symmetry searches, other parameters unchanged. The best class (163,123 segments) underwent 3D auto-refinement using Blush regularization with the AmyBlush (amy-v1.0) network in RELION v5.1. The other parameters changed from the defaults were as follows: angular sampling interval = 1.8°, outer tube diameter = 175 Å, number of unique asymmetrical units = 3, initial twist = −0.11°, initial rise = 4.8 Å, central Z length = 20%, min twist = −0.2°, max twist = 0°, twist step = 0.01°, min rise = 4.75 Å, max rise = 4.85 Å, rise step = 0.01 Å. The refined map underwent mask creation (initial binarization threshold = 0.001, binary map extension and soft edge = 10 pixels, central Z length = 20%) and standard post-processing to give the final PDD Lewy aSyn map at an estimated global resolution of 2.3 Å (Supplementary Fig. S9).

### Model building, refinement, and analysis

Initial model building was performed by fitting the existing high-twist aSyn Lewy fold (PDB: 8A9L) into the refined maps. Subsequent model fitting and refinement was performed using ISOLDE[5]. After fitting into the density, each main protofilament was translated to give a stack of 3 strands. After Ramachandran parameters, rotamers, and clashes were satisfied in the middle strand, the ISOLDE command “isolde write phenixRsrInput #

<MODEL><MAP resolution> #

<MAP>” was used to export the model and generate a rigid-body refinement settings file for real-space refinement in PHENIX[1].

The outer 2 rungs were deleted, and unknown peptides in island A and B were rigid-body fit. For DLB, only island B was fit. The models were then translated into a stack of 9 rungs. Data were deposited into the Protein Data Bank (PDB), under accession codes 38GH (DLB) and 38GI (PDD), and into the Electron Microscopy Data Bank (EMDB), under accession codes EMD-78792 (DLB) and EMD-78793 (PDD).

Amyloid packing difference (APD) scores were calculated and associated figures were generated as previously described[25]. Other structural figures were made using University of California San Francisco (UCSF) ChimeraX[21].

## RESULTS

We partially purified sarkosyl-insoluble filaments from the postmortem brain of a 74-year-old female patient who carried a clinical diagnosis of DLB and was found to have underlying diffuse neocortical LBD (Parkinson’s disease Braak Stage 6) (Fig. 1a, Supplementary Fig. S1), keeping a small amount of material at each step for analysis (denoted as fractions 1–5) (Supplementary Fig. S1). Negative-stain electron microscopy of the final purified fraction confirmed the presence of filaments (Supplementary Fig. S2). To assess the pathogenicity of these filaments, we used each purification fraction (fractions 1–5) to transfect a stable human embryonic kidney 293T (HEK293T) cell line previously shown to specifically respond to Lewy prions[4]. These cells express aSyn 35–110 with the E46K mutation fused with yellow fluorescent protein (YFP) tag at the C-terminus (aSyn(35–110)*E46K-YFP). Transfecting cells with increasing seed concentrations revealed a robust dose-response relationship of infectivity, measured as percentage of cells with aggregates (CWA%) (Supplementary Fig. S10).

**Fig. 1.**
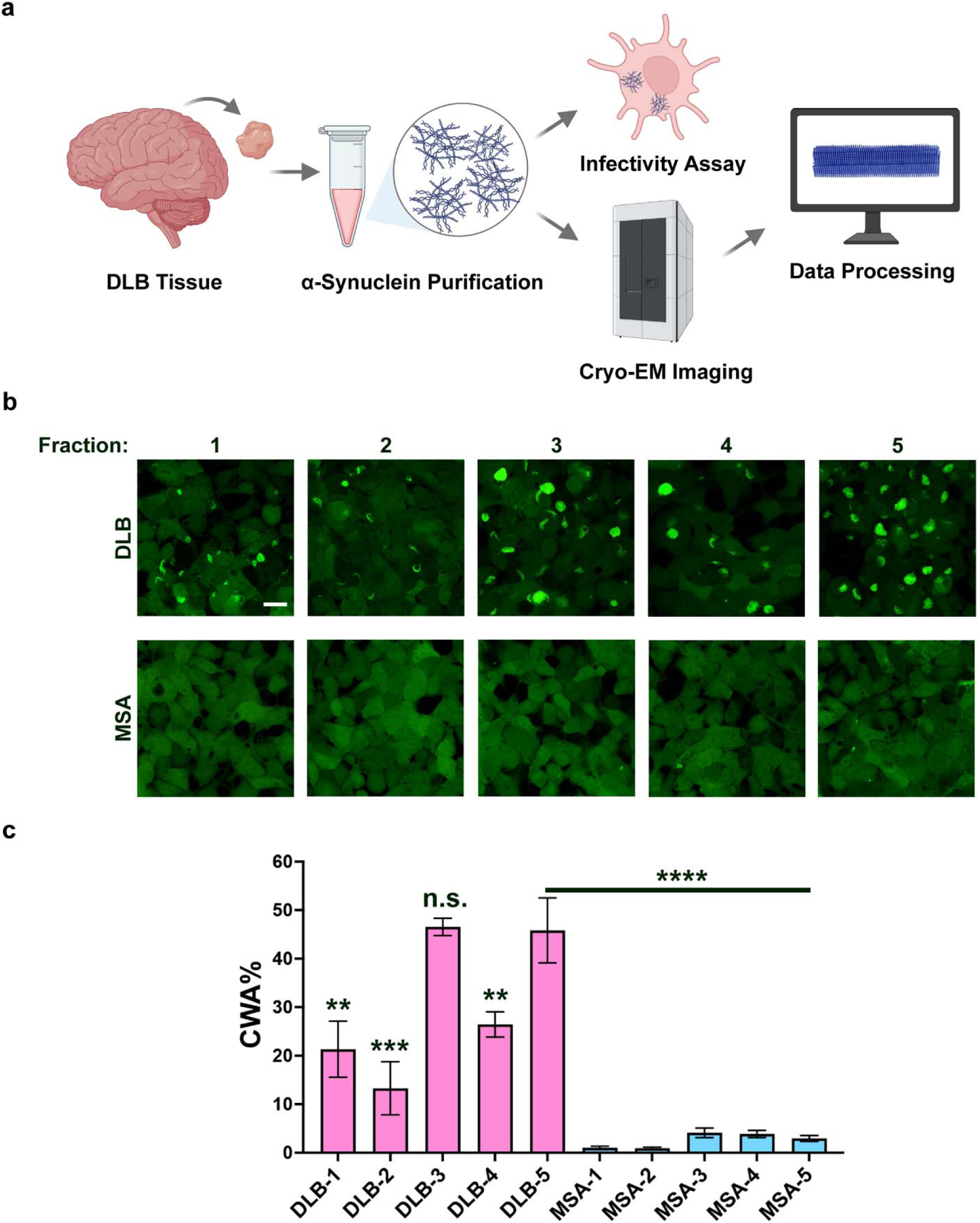
Cell infectivity of DLB and MSA brain fractions in aSyn(35–110)*E46K-YFP cells. **a** Summary of sample preparation and data acquisition in this study. **b** Representative images of cells infected with different DLB or MSA fractions (1–5). YFP is shown in green. Scale bar is 20 μm. **c** Bar graph showing aSyn(35–110)*E46K-YFP cells with aggregates (CWA%), plotted as a percent of total number of cells. To calculate the cell number, nuclei were stained with Hoechst (not shown here). The infectivity of DLB fractions and MSA fractions is depicted by pink bars and blue bars, respectively. Data shown as mean ± SD. Unpaired Student’s *t*-test was performed to compare infectivity of individual brain fractions against their respective fraction 5. Student’s *t*-test was also performed to compare infectivity of DLB fraction 5 and MSA fraction 5. \*\**P* < 0.01, \*\*\**P* < 0.001, \*\*\*\**P* < 0.0001; n.s., non-significant.

When comparing infectivity across the fractions at the highest seed concentration (1,562.5 pg total protein/μL), we observed visibly larger intracellular aggregates and maximum CWA% in fractions 3 and 5 (Fig. 1b, c). Importantly, this construct specifically blocks MSA prions by inhibiting the formation of the structurally critical E46-K80 salt bridge in these aSyn folds[2, 4], enabling discrimination of MSA vs. LB prions in the inocula. Significantly lower infectivity of fractions prepared in the same way from the brain tissue of a patient with MSA (Fig. 1c, see Methods) confirmed that this cellular aggregation is in response to Lewy prions specifically (full dataset in Supplementary Fig. S10). As an additional control to ascertain that the MSA fractions were infectious, we generated another HEK293T cell line that stably expressed the A53T variant of aSyn (35–110) with a C-terminal YFP tag, similar to one that was previously engineered to facilitate propagation of both MSA and LB prions[4]. We detected prominent intracellular aggregation of A53T aSyn in response to both MSA and DLB fractions. However, these constructs propagated MSA and LB prions more similarly to each other (Supplementary Fig. S11) than the previously generated cell line[4]12303}, perhaps owing to differences in the cell lines or patient samples. Despite this, we were able to confirm the selectivity of our E46K cell line.

We next performed cryo-EM imaging on DLB fraction 5 (Fig. 2a), which was the most infectious in our aSyn(35–110)*E46K-YFP cell line and was confirmed to contain filaments via negative-stain EM (Supplementary Fig. S2). Subsequent reference-free 2D classification revealed that most of the class averages (∼70% of segments) lacked discernable twist, a minority (∼20%) exhibited a very subtle twist without visible crossover, and the remaining segments were low-resolution particles (Fig. 2b, c, Supplementary Figs. S4–S5). We generated an initial model *de novo* using the twisted classes and then, using both twisted and untwisted segments, performed iterative 3D classification and helical refinement. This resulted in a 2.9-Å reconstruction of low-twist aSyn filaments with a crossover distance of 7,500 Å (Fig. 2d, e, Supplementary Table S1). Blush regularization, which uses trained data to enhance reconstructions of low signal-to-noise particles[17, 20], was critical to reduce overfitting and thus enabled high-resolution reconstructions of these low-twist filaments (see Methods and Supplementary Figs. S3, S6, S7, S9, S12).

**Fig. 2.**
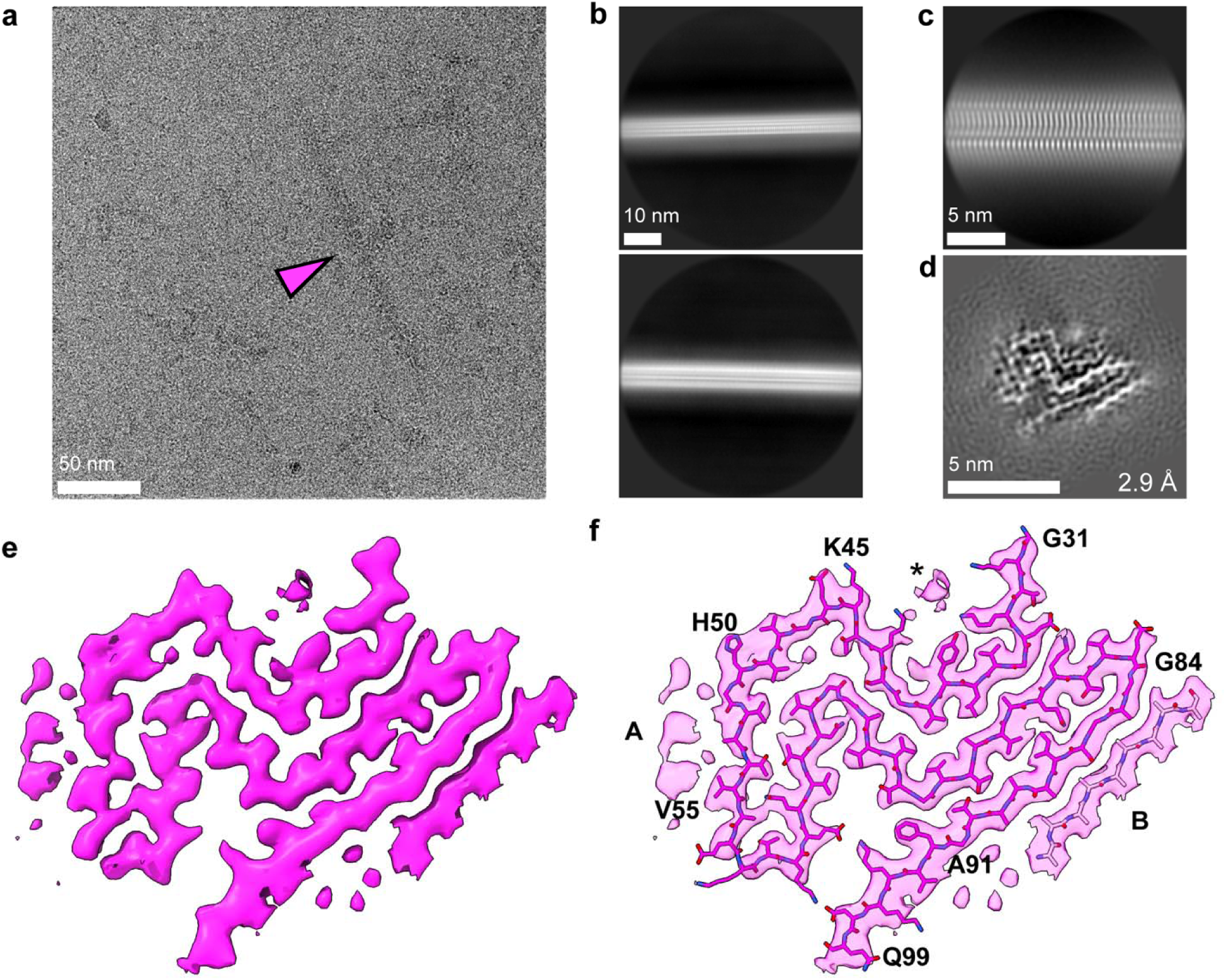
Cryo-EM structures of low-twist aSyn filaments from a patient with DLB. **a** Example micrograph of an aSyn filament labeled with an arrow. **b** Representative filament 2D class averages at 900 pixels (binned to 300 pixels; box size, ∼655 Å). **c** Representative filament 2D class average at 288 pixels (∼210 Å). **d** Cross-section of aSyn filament reconstruction perpendicular to the helical axis. **e** Cross-section of aSyn filament density map showing main chain density and surrounding peptidic and non-peptidic densities. **f** Model of long-crossover Lewy aSyn fitted into the map. Residues in the termini of the ordered core are labeled, along with residues delineating the regions where disconnected unidentified densities are observed. A and B are 2 disconnected peptidic densities. *Location of N-terminal cofactor.

We resolved residues 31–99 of aSyn, which adopt a 3-layered fold that is extremely similar to the canonical Lewy fold (Fig. 2f). The map contained patterns of additional density that were disconnected from the main protofilament observed in the previously reported Lewy aSyn map. The N-terminal Lys-rich cavity contained density consistent with a bound cofactor (previously proposed to be polyphosphate[14]), as well as a putative ordered water molecule between Y39 and K43. Two peptidic densities were also observed: a weaker ∼6-residue “island A” adjacent to His50-Val55 in the hinge region and a more well-resolved 9-residue “island B” associated with G84-A91 of the C-terminal hairpin. The density for island A was insufficient to model the backbone, and attempts to identify island B using ModelAngelo and the wild-type (WT) aSyn sequence as an input were unsuccessful[15].

Comparison of our low-twist structure with high-twist Lewy filaments (PDB: 8A9L) revealed 2 main differences. First, the low-twist filaments were left-handed, in contrast to the right-handed high-twist filaments (Fig. 3a). The handedness was confirmed by the fit of carbonyl oxygens in the polypeptide chain to the density. Structural alignment showed that the folds were highly similar in the XY plane (perpendicular to the filament Z-axis), with a root-mean-square deviation (RMSD) of 2.3 Å. However, visualizing the folds along the Z-axis demonstrated that the low-twist Lewy fold adopted a “flatter” conformation compared to the high-twist fold (Fig. 3b). While the C-terminal hairpins were superimposable, both the Z-registers of the hinge region and the N-terminal cavity were shifted to be more like the register of the hairpin. The maximum displacement between the low- and high-twist forms was 3.1 Å in the hinge region (K58 Cα) and 4.8 Å in the N-terminal cavity (G31 Cα). Because of the shift in register, the bridging interactions between the N-terminal cavity and C-terminal hairpin, such as the salt bridges between E35 and K80, were displaced by one protofilament rung but were otherwise intact (Fig. 3c). The similarities between high-twist and low-twist Lewy protofilaments were reaffirmed by the low amyloid packing difference (APD) scores (4% and 16% in XY and XYZ, respectively; Fig. 3d)[25]. We did note one contact (K60-K96) that was only present in the high-twist PF but attributed this to modeling stochasticity due to incomplete density of the K60 sidechain.

**Fig. 3.**
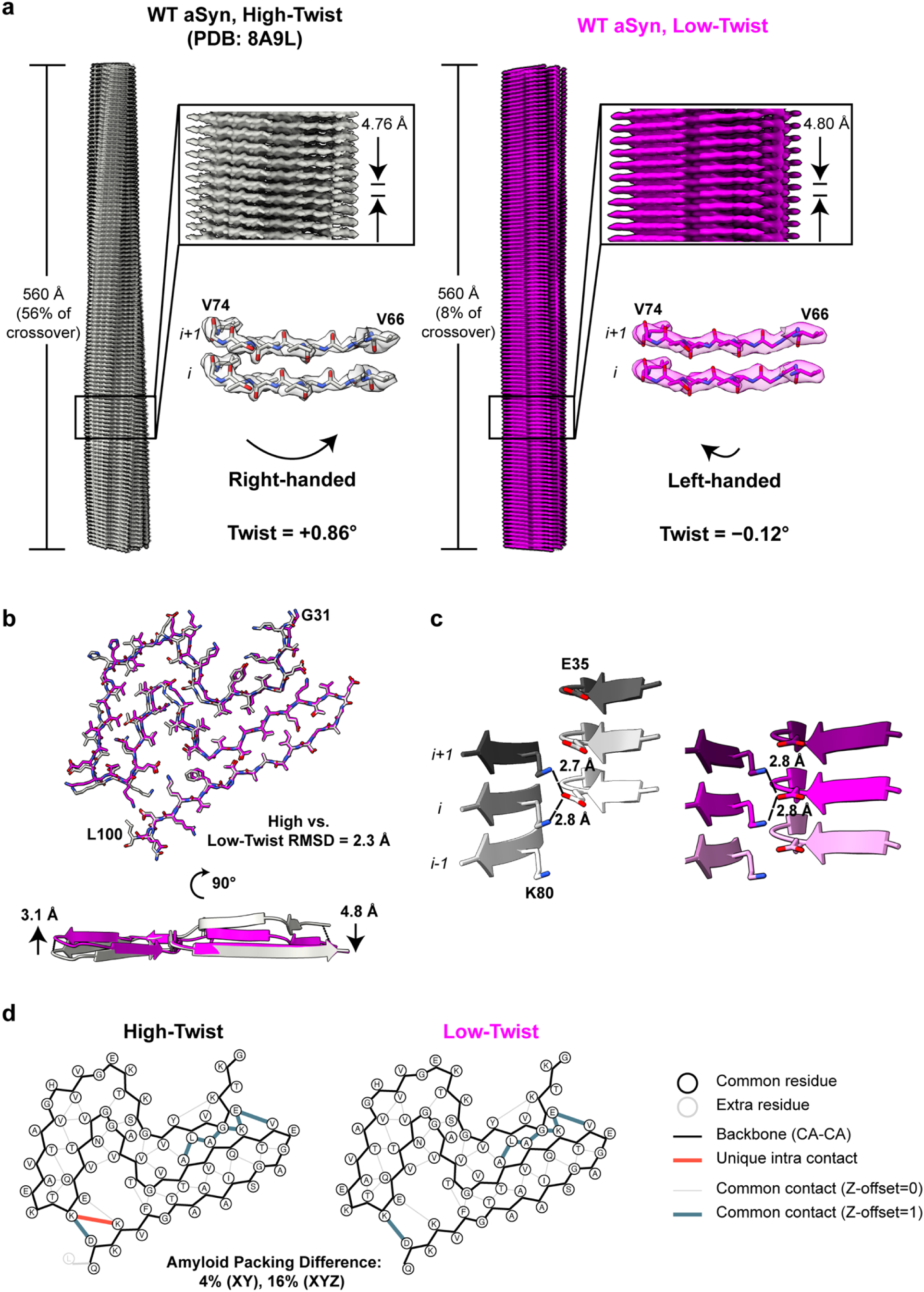
Comparison of high-twist (PDB: 8A9L) and low-twist Lewy aSyn filaments. **a** Helical symmetry parameters and handedness of high-twist (gray) and low-twist (purple) filaments. Handedness assignments are confirmed by the fit of backbone carbonyl oxygens into the density. **b** Superposition of low-twist and high-twist aSyn models, showing both a cross-sectional all-atom view and a ribbon view along the filament axis. **c** Comparison of interstrand salt bridges between Lys80 and Glu35 in high-twist and low-twist filaments. **d** Amyloid packing differences between high-twist and low-twist filaments.

Encouraged by the ability to resolve low-twist filaments to high resolution, we next wanted to assess whether the differences in structure were based on filament selection or patient-to-patient variability. To this end, we revisited a publicly available dataset (EMPIAR-11140, from an 87-year-old male with PDD; PDD stage and Parkinson’s disease Braak stage were not reported) deposited by Yang *et al*. as part of their 2022 study, wherein the authors obtained reconstructions of high-twist filaments[33]. Low-twist or non-twisted 2D averages make up the majority of aSyn filaments from this patient, but these were excluded from further 3D reconstruction. We extracted all filament segments from the provided start-end coordinates (Supplementary Fig. S8) and used a low-pass-filtered version of the DLB map as a reference for 3D classification of this dataset. Subsequent processing enabled by Blush regularization yielded a left-handed 2.3-Å Lewy aSyn reconstruction with a shorter crossover distance (5,600 Å; Fig. 4a) compared to our low-twist DLB reconstruction and extremely similar patterns of additional unidentified densities. The models were nearly identical, with a RMSD of 0.35 Å between the α-carbons of the protofilaments (Fig. 4b, c). These results indicate that low-twist Lewy aSyn filaments are structurally conserved between different patients with Lewy pathology.

**Fig. 4.**
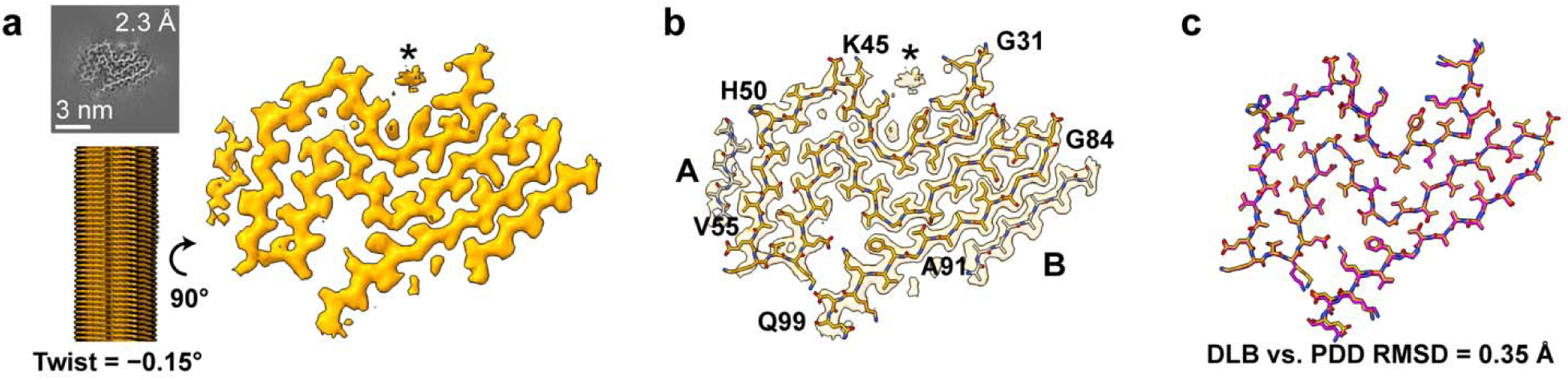
Cryo-EM structures of low-twist aSyn filaments from a patient with PDD (EMPIAR-11140). **a** Reconstruction of low-twist filament showing both a side-on view and a one-rung cross-section. **b** Model of long-crossover Lewy aSyn fitted into the map. Residues in the termini of the ordered core are labeled, along with residues delineating the regions where disconnected unidentified densities are observed. Two disconnected peptidic densities are seen: A and B. Both density islands are modeled as polyalanine. **c** Main protofilament alignment between long-crossover DLB aSyn (purple) and long-crossover PDD aSyn (orange). *Location of N-terminal cofactor.

## DISCUSSION

Our results demonstrate that aSyn filaments isolated from the brain of a patient who presented with DLB are infectious and can undergo templated propagation in a cell line expressing truncated (residues 35–110), YFP-tagged E46K aSyn, which does not propagate MSA prions. The filaments purified from brain tissue of a patient with DLB consisted predominately of low-twist aSyn with the same general architecture as the canonical Lewy fold. The protofilament core contained residues 31 to 99, so it is notable that robust propagation in cells can occur even in the absence of 4 N-terminal residues that flank an unknown cofactor. In the Lewy fold, E46 is a surface-exposed residue near the N-terminal cationic pocket and may form a salt bridge with the adjacent surface-exposed K45 (Fig. 2 and Supplementary Fig. S13). However, in all MSA protofilament types, E46 forms a salt bridge with K80. This bridging of distant regions in MSA protofilaments is postulated to contribute significantly to protofilament stability[2, 26, 31]. These structural differences help explain the ability of E46K aSyn to propagate the Lewy fold specifically. To further validate the ability of this cellular system to propagate Lewy aSyn, future studies will need to investigate the structures of filaments isolated from these cells and test additional aSyn constructs to understand the requirements for Lewy fold propagation.

We also demonstrate that Lewy aSyn filaments adopt a conserved structural core despite substantial differences in twist to previously reported structures. Thus, we provide a more complete high-resolution picture of pathologically relevant aSyn filament types in patients with distinct clinical presentations of LBD. Low-twist filaments, which constitute most filaments isolated from patients with LBs, have historically been refractory to high-resolution cryo-EM structural determination. Here, we show that Blush regularization greatly enhances reconstructions of these filaments. This approach may be broadly applicable to other low-twist or apparently straight filaments.

Recent work by Zhang *et al*. reported structures of *ex vivo* aSyn filaments from patients with LBD carrying single point mutations (A53T, G51D, H50Q)[34]. Notably, low-twist A53T and G51D aSyn exhibited crossover distances similar to those observed in our WT structures. In the structures reported by Zhang *et al*., the core Lewy fold is intact, which is further corroborated by the infectivity of the DLB *ex vivo* aSyn filaments in an A53T cell line. The mutant aSyn filaments lacked density for island A near the hinge region and frequently formed doublet filaments (exclusively in the case of G51D), likely reflecting steric or electrostatic effects of these hinge region mutations. In contrast, our WT low-twist structures retained island A density, consistent with high-twist WT and H50Q filaments. Low-twist A53T and G51D filaments are also left-handed, like our WT low-twist structures, whereas high-twist WT and H50Q filaments are right-handed.

Given that mutant aSyn accounts for <1% of LB cases[18, 23], our structures of low-twist WT aSyn link previously described high-twist WT structures with low-twist mutant aSyn.

Finally, although the overall fold is conserved, differences in helical twist alter the filament surface and may influence interactions with small molecules or antibodies that engage multiple filament strands simultaneously. Thus, high-resolution structures of low-twist aSyn filaments may provide an important framework for the design of probes or therapeutics targeting Lewy body pathology.

## CONTRIBUTIONS

NLY, VGC-V, and GEM conceived the study. MM purified patient tissue. NLY, MM, AAM, and ET contributed to cryo-EM data collection. NLY processed cryo-EM data, generated atomic models, and performed structural analysis. VGC-V performed infectivity experiments and analyzed infectivity data. WWS and DAM contributed to the neuropathological diagnoses. NLY, MM, VGC-V, and GEM drafted the manuscript. NLY, MM, AAM, JIA, DAM, WWS, SBP, ET, VGC-V, and GEM revised the manuscript and approved the final version.

## ETHICS DECLARATION

### Consent for Publication

Not applicable. No individual-level data reported.

### Data Availability

Cryo-EM maps and atomic coordinates have been deposited in the EMDB and PDB with the following accession codes: EMD-WWWWW (DLB), EMD-XXXXX (PDD), and PDB YYYY (DLB), PDB ZZZZ (PDD). Any other relevant data are available from the corresponding author upon request.

## ACKNOWLEDGEMENTS

Brain tissue from the patient with DLB was provided by the UCSF Neurodegenerative Disease Brain Bank. Brain tissue from the patient with MSA was provided by the Massachusetts Alzheimer’s Disease Research Center. The authors acknowledge the invaluable contributions of the study participants and families as well as the assistance of the support staffs at each of the participating sites. Portions of Figure 1 and Supplementary Figure S1 were made using BioRender.com. The authors also thank Dr. Katie Hatstat for her critical feedback on this manuscript.

