## Supplementary Information for "Cryo-EM Structures of Infectious Low-Twist α-Synuclein Filaments in Lewy Bodies"

<sup>1</sup>Institute for Neurodegenerative Diseases, University of California San Francisco, San Francisco, CA, USA; <sup>2</sup>Department of Neurology, University of California San Francisco, San Francisco, CA, USA; <sup>3</sup>Department of Pathology, University of California San Francisco, San Francisco, CA, USA; <sup>4</sup>Weill Institute for Neurosciences, Memory and Aging Center, Department of Neurology, University of California San Francisco, San Francisco, CA, USA; <sup>5</sup>Department of Biochemistry and Biophysics, University of California San Francisco, San Francisco, CA, USA; <sup>6</sup>Department of Pharmaceutical Chemistry, University of California San Francisco, San Francisco, CA, USA

\*Correspondence: Gregory E. Merz

**Table S1. Cryo-EM data collection and refinement statistics.**

|  | **DLB** | **PDD** |
| --- | --- | --- |
| **PDB accession code** | **YYYY** | **ZZZZ** |
| **EMDB accession code** | **EMD-XXXXXX** | **EMD-XXXXXX** |
| **Data collection and processing** |  |  |
| Microscope and camera | Krios G4, Falcon 4i | Dataset downloaded from EMPIAR-11140 |
| Nominal magnification | 165,000× |  |
| Voltage (kV) | 300 |  |
| Data acquisition software | EPU |  |
| Exposure navigation | Image shift |  |
| Electron exposure (e<sup>-</sup>/Å<sup>2</sup>) | 45 |  |
| Defocus range (μm) | -0.8 to -1.0 |  |
| Pixel size (Å) | 0.728 | 0.727 |
| Symmetry imposed | C1 | C1 |
| Initial segments (no.) | 331,207 | 725,340 |
| Final segments (no.) | 60,550 | 163,123 |
| Map resolution (Å) | 2.9 | 2.3 |
| FSC threshold | 0.143 | 0.143 |
| Helical twist | -0.12 | -0.15 |
| Helical rise (Å) | 4.80 | 4.79 |
| **Refinement** |  |  |
| Model resolution (Å) | 3.2 | 2.5 |
| FSC threshold | 0.5 | 0.5 |
| Map sharpening B factor (Å<sup>2</sup>) | -69.4 | -51.1 |
| **Model composition** |  |  |
| Non-hydrogen atoms | 4,734 | 4,995 |
| Protein residues | 621 | 621 |
| **B factors (Å<sup>2</sup>)** |  |  |
| Protein | 83.9 | 43.7 |
| **R.M.S. deviations** |  |  |
| Bond lengths (Å) | 0.008 | 0.008 |
| Bond angles (°) | 1.077 | 1.087 |
| **Validation** |  |  |
| MolProbity score | 1.04 | 0.76 |
| Clash score | 2.56 | 0.82 |
| Rotamer outliers (%) | 0.00 | 0.00 |
| **Ramachandran plot** |  |  |
| Favored (%) | 98.51 | 100.00 |
| Allowed (%) | 1.49 | 0.00 |
| Outliers (%) | 0.00 | 0.00 |

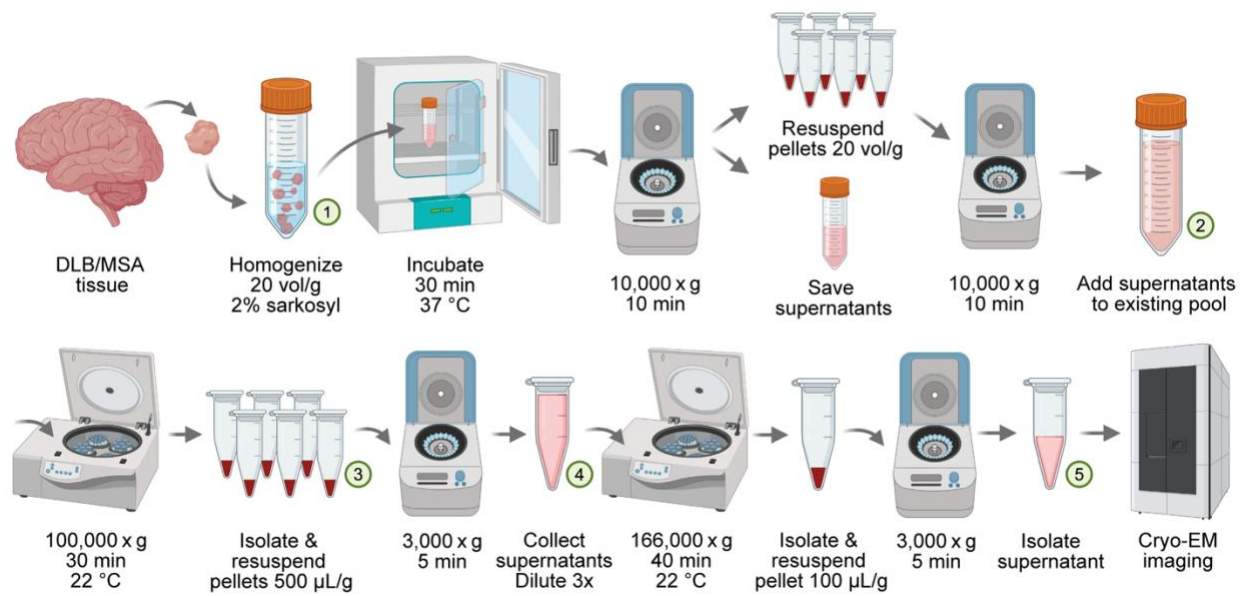

**Fig. S1** Detailed flowchart of filament sample preparation from DLB and MSA brain tissue. Fractions used in the cell infectivity assay are numbered 1 to 5, indicating the point of collection.

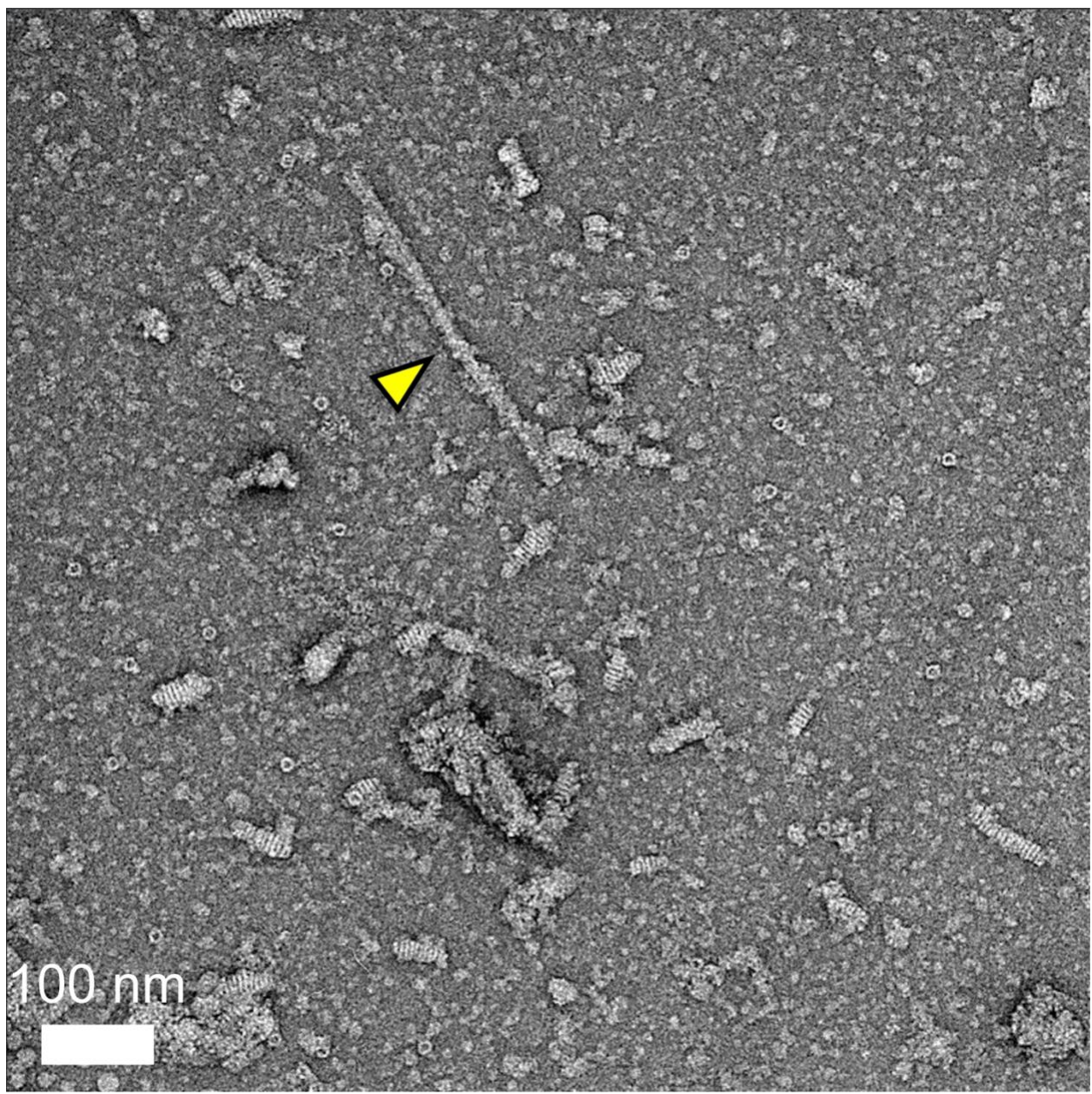

**Fig. S2** Representative negative-stain transmission electron micrograph of purified DLB filaments (yellow arrow).

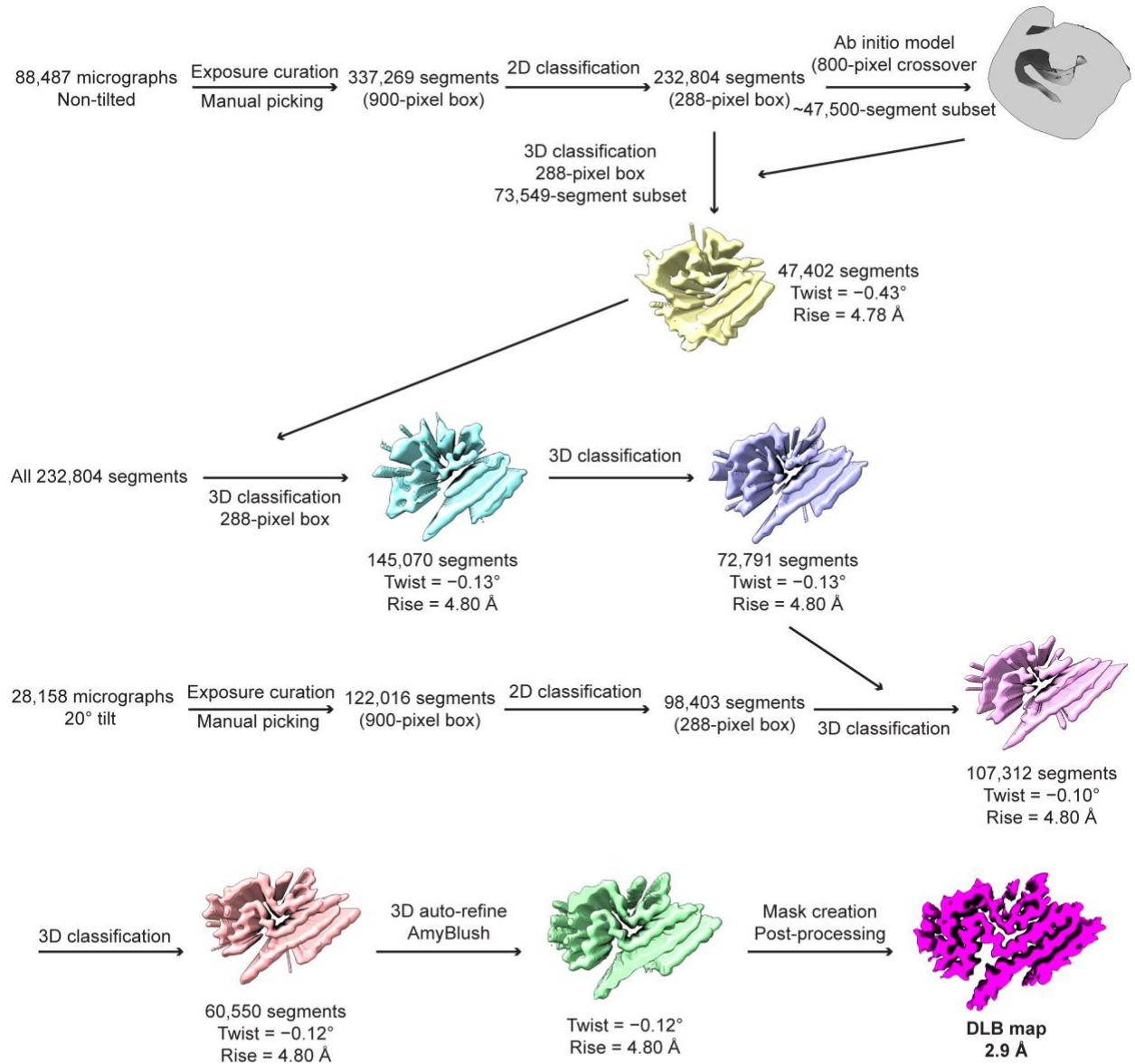

**Fig. S3** Data processing flowchart for the DLB dataset, which was collected in house.

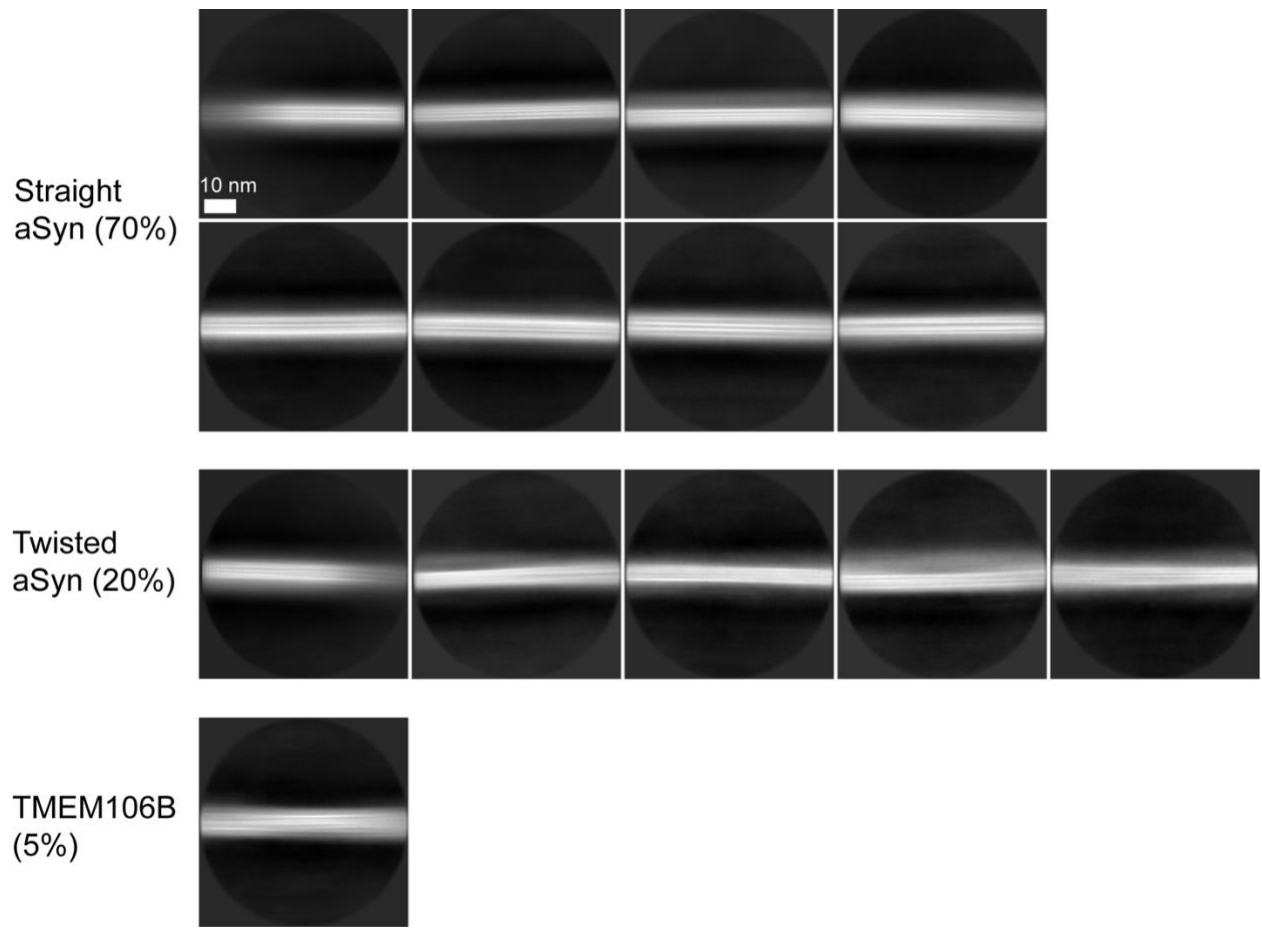

**Fig. S4** Reference-free 2D class averages of filaments from non-tilted DLB datasets (900 pixels binned to 300 pixels; final Å/pixel of 2.184) after 15 iterations. Scale bar represents 10 nm. The twisted classes were used for *ab initio* model generation. However, both the straight and twisted aSyn filaments were pooled and used for 3D classification due to the low number of segments (47,000) in the twisted classes. Note that the twisted aSyn filaments tend to converge into straight filament classes if run for more iterations.

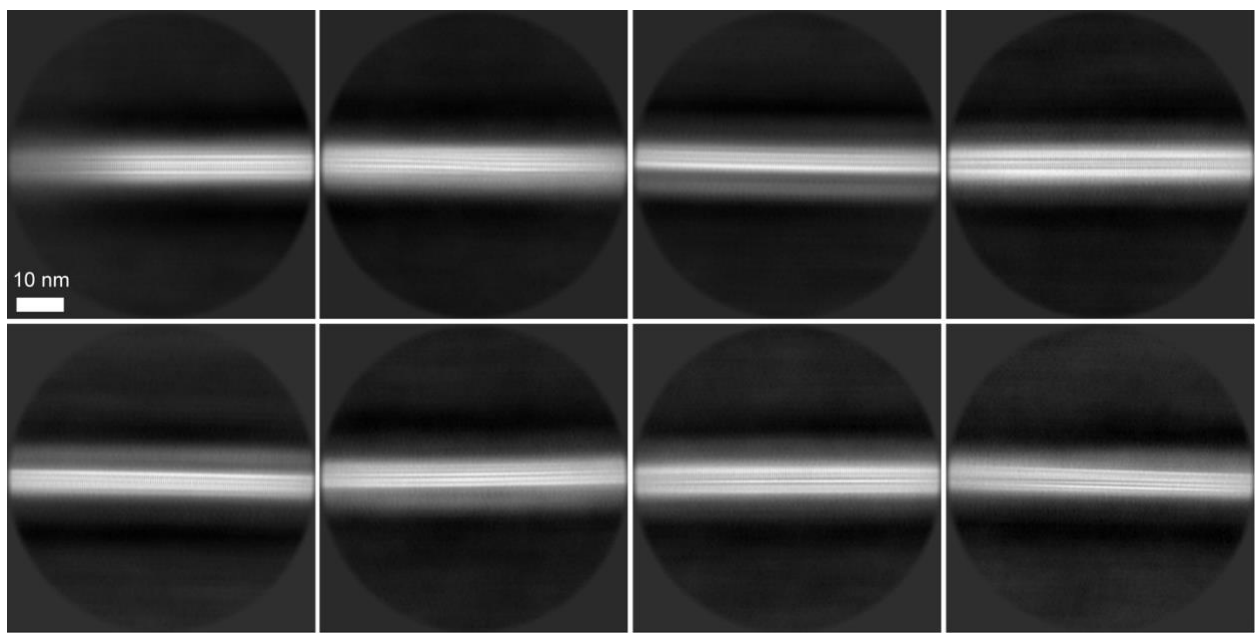

**Fig. S5** Reference-free 2D class averages of filaments from tilted DLB dataset (900 pixels binned to 300 pixels; final Å/pixel of 2.184) after 25 iterations. Scale bar represents 10 nm.

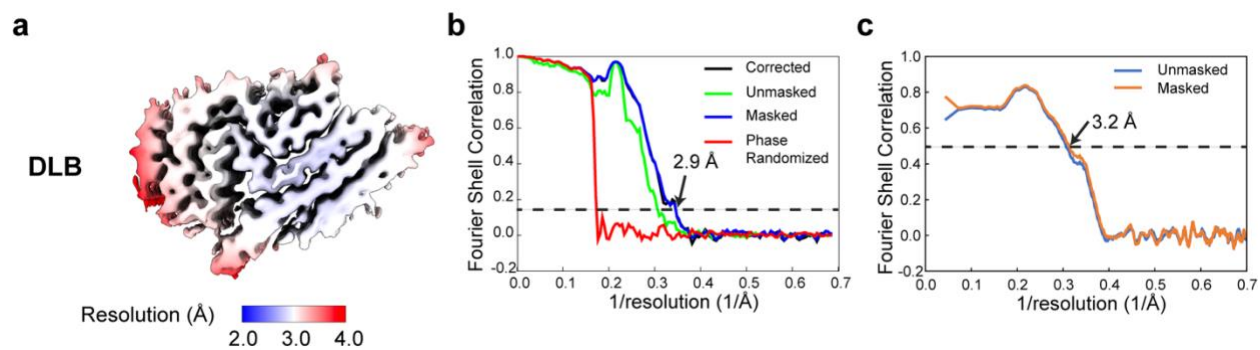

**Fig. S6** Resolution estimates for the DLB map and model. **a** Local resolution map of DLB map. **b** Fourier Shell Correlation (FSC) curves of refined DLB half-maps. **c** Map-to-model FSC curves of DLB.

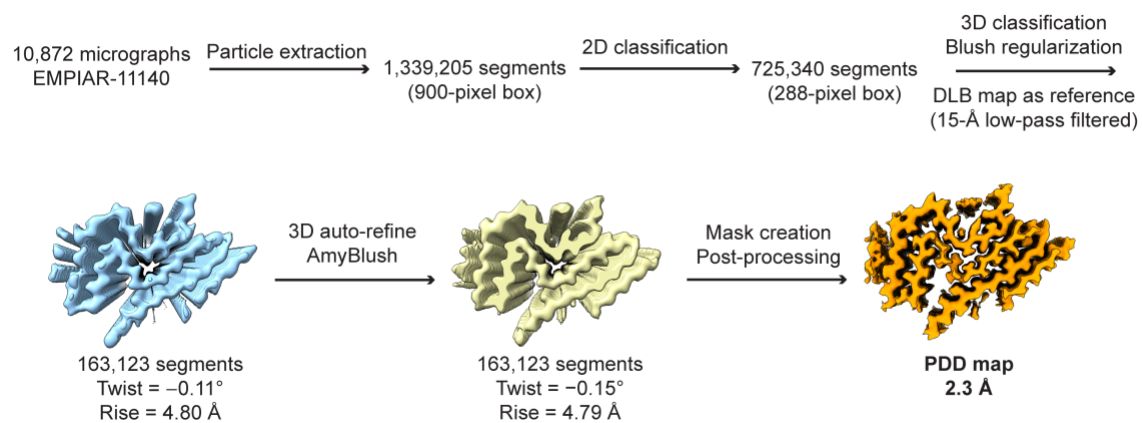

**Fig. S7** Data processing flowchart for the PDD dataset, which was downloaded from EMPIAR.

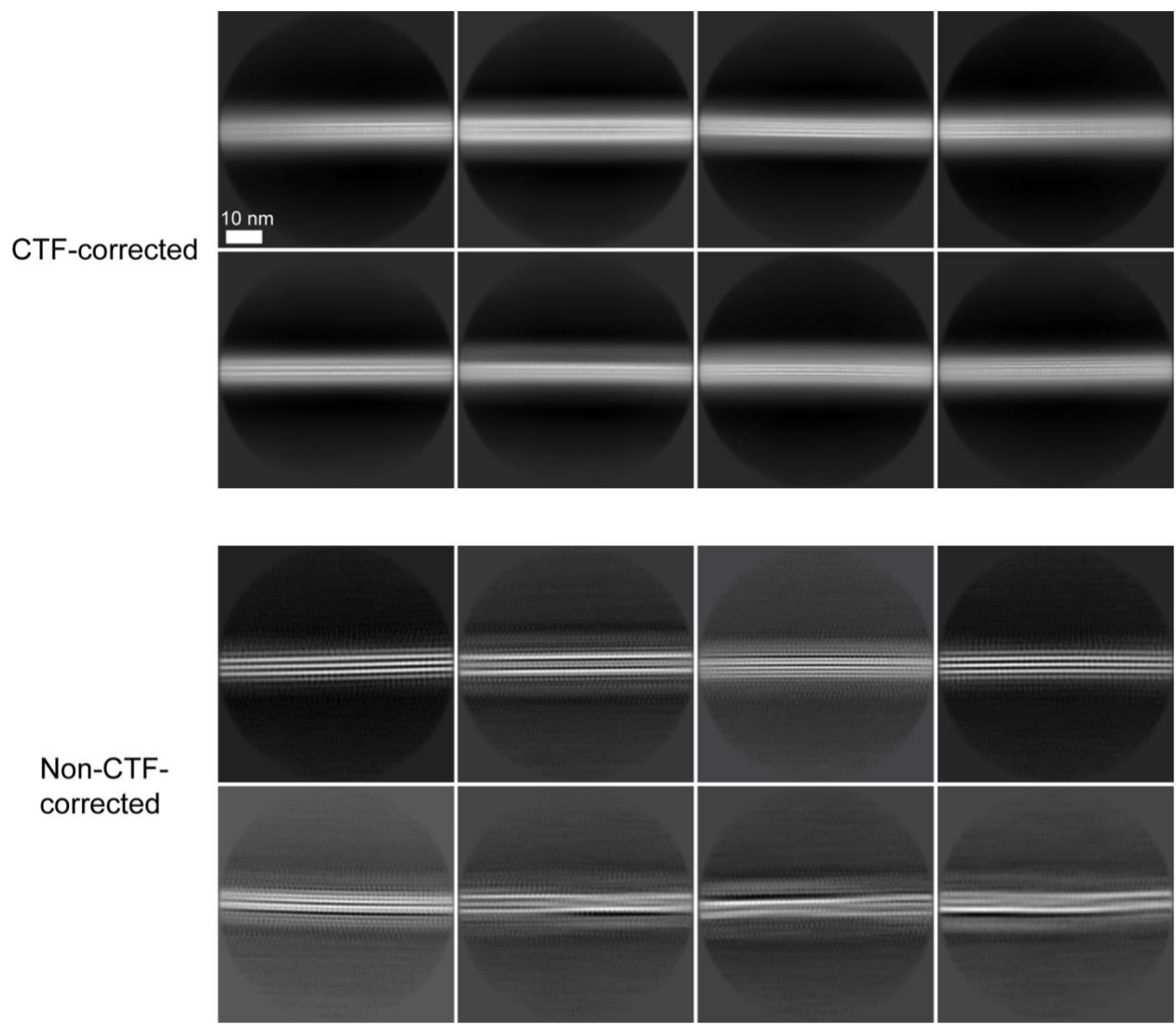

**Fig. S8** Reference-free 2D class averages of filaments from PDD dataset (EMPIAR-11140; 900 pixels binned to 300 pixels; final Å/pixel of 2.181) after 25 iterations. Scale bar represents 10 nm. The top panels show CTF-corrected 2D averages, showing clear β-sheet spacing. To examine potential overfitting, we also obtained non-CTF-corrected averages, which better separated out twisted classes but resulted in lower resolution.

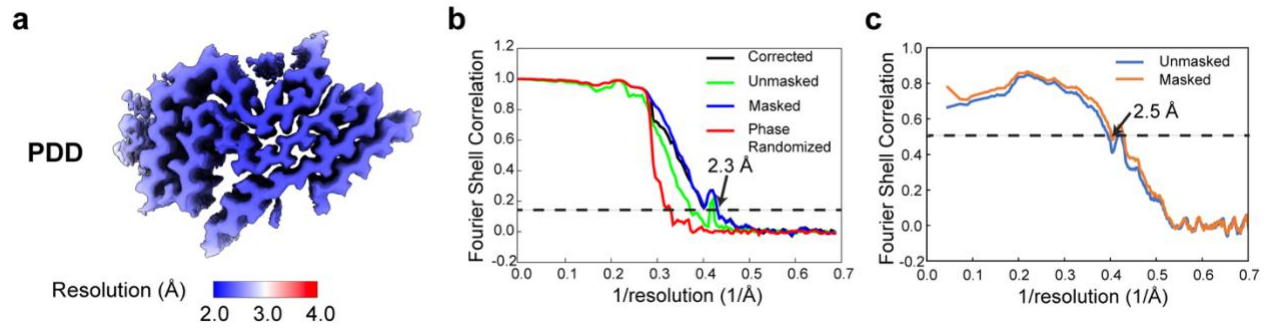

**Fig. S9** Resolution estimates for the PDD map and model. **a** Local resolution map of PDD map. **b** FSC curves of refined PDD half-maps. **c** Map-to-model FSC curves of PDD.

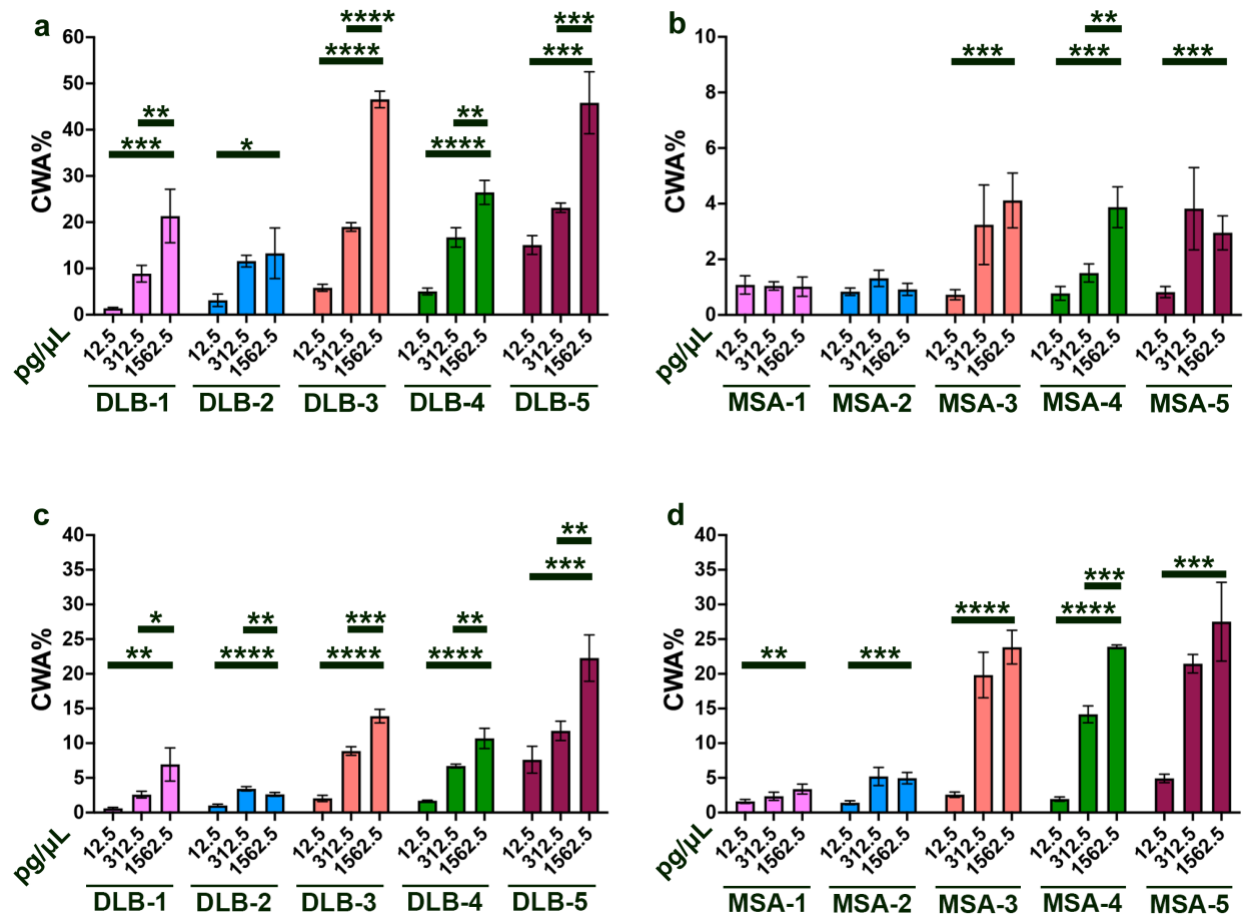

**Fig. S10** Cellular infectivity of brain fractions dictated by the seed concentration. **a, b** CWA% of aSyn(35–110)\*E46K-YFP cells infected with different brain fractions of either DLB (**a**) or MSA (**b**) sample. Cellular transfections were performed with the following increasing dose of brain fractions (total protein concentration): 12.5 pg/μL, 312.5 pg/μL, or 1,562.5 pg/μL. Total number of cells were calculated with Hoechst nuclear stain. **c, d** CWA% of aSyn(35–110)\*A53T-YFP cells infected with either DLB (**c**) or MSA (**d**) brain fractions. Transfections and cellular count were performed as described above. Data shown as mean ± SD. Unpaired Student's *t*-test performed for comparing infectivity of different seed concentrations within individual fractions. \**P* < 0.05, \*\**P* < 0.01, \*\*\**P* < 0.001, \*\*\*\**P* < 0.0001.

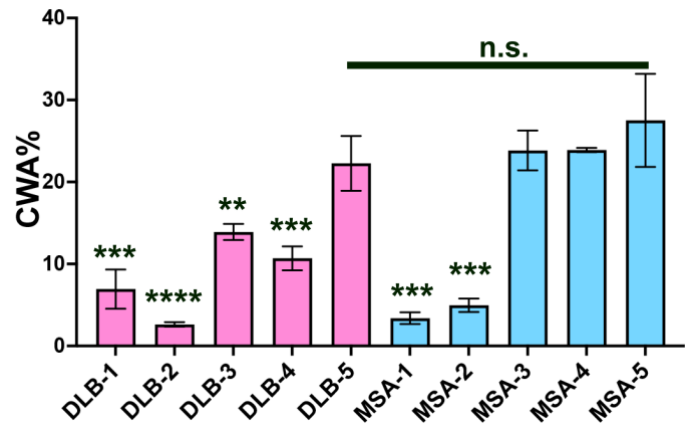

**Fig. S11** Infectivity of DLB and MSA brain fractions in aSyn(35–110)\*A53T-YFP cells. CWA% of aSyn(35–110)\*A53T-YFP cells infected with different brain fractions. Total cell numbers were calculated based on Hoechst nuclear stain. Graphs show data for DLB fractions in pink columns and MSA fractions in blue columns. Data shown as mean ± SD. Using unpaired Student's *t*-test, infectivity was compared between individual brain fractions and fraction 5 of the respective groups. The same statistical tests were conducted between fractions 5 of DLB and MSA brains. \*\*$P < 0.01$, \*\*\*$P < 0.001$, \*\*\*\*$P < 0.0001$.

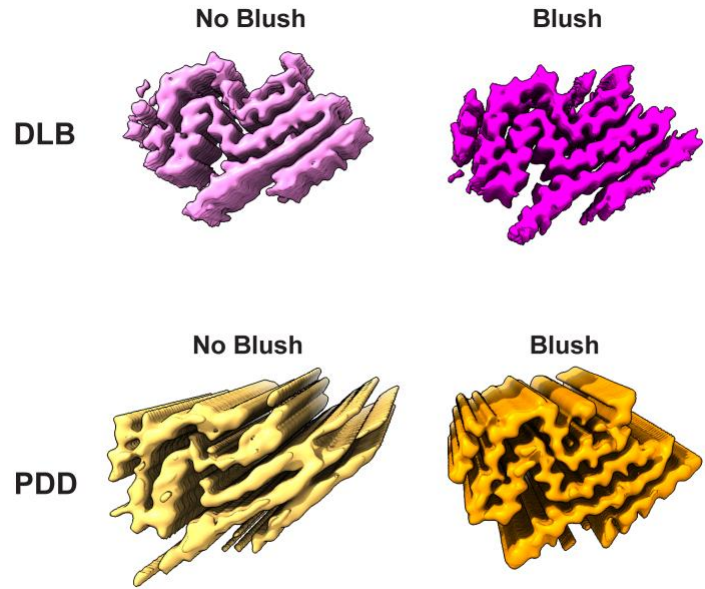

**Fig. S12** Blush regularization during 3D classification or auto-refinement enhances low-twist Lewy aSyn reconstructions. The DLB comparisons are after multiple rounds of 3D classification followed by refinement, mask creation, and post-processing. Blush regularization (top right) was applied during auto-refinement. The PDD comparisons are after one round of 3D classification, in which the appearance of streaky density characteristic of overfitting is prevented by Blush regularization.

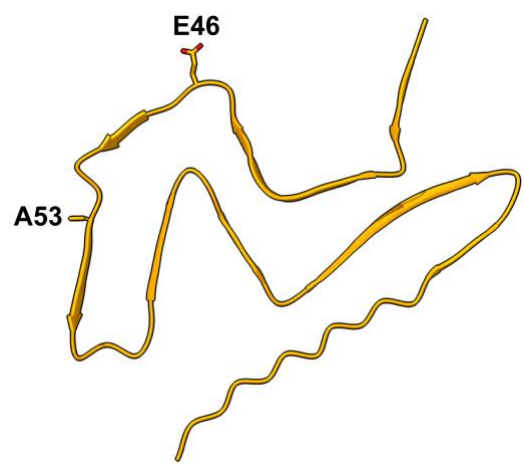

**Fig. S13** Locations of the Ala53 and Glu46 residues in the Lewy αSyn fold.
